# Principles governing the topological organization of object selectivities in ventral temporal cortex

**DOI:** 10.1101/2021.09.15.460220

**Authors:** Yiyuan Zhang, Ke Zhou, Pinglei Bao, Jia Liu

## Abstract

To achieve the computational goal of rapidly recognizing miscellaneous objects in the environment despite large variations in their appearance, our mind represents objects in a high-dimensional object space to provide separable category information and enable the extraction of different kinds of information necessary for various levels of the visual processing. To implement this abstract and complex object space, the ventral temporal cortex (VTC) develops different object-selective regions with certain topological organization as the physical substrate. However, the principle that governs the topological organization of object selectivities in the VTC remains unclear. Here, equipped with the wiring cost minimization principle constrained by the wiring length of neurons in human temporal lobe, we constructed a hybrid self-organizing map (SOM) model as an artificial VTC (VTC-SOM) to explain how the abstract and complex object space is faithfully implemented in the brain. In two *in silico* experiments with the empirical brain imaging and single-unit data, our VTC-SOM predicted the topological structure of fine-scale functional regions (face-, object-, body-, and place-selective regions) and the boundary (i.e., middle Fusiform Sulcus) in large-scale abstract functional maps (animate vs. inanimate, real-word large-size vs. small-size, central vs. peripheral), with no significant loss in functionality (e.g., categorical selectivity, a hierarchy of view-invariant representations). These findings illustrated that the simple principle utilized in our model, rather than multiple hypotheses such as temporal associations, conceptual knowledge, and computational demands together, was apparently sufficient to determine the topological organization of object-selectivities in the VTC. In this way, the high-dimensional object space is implemented in a two-dimensional cortical surface of the brain faithfully.

## Introduction

The natural world we live in includes various kinds of objects. To make sense of and interact with the world, we need to identify, discriminate, and categorize these objects into groups based on their characteristics. How is this diversity of objects represented in mind? It is suggested that objects are represented metaphorically as points in a high-dimensional object space in our minds (Hebart, Zheng, Pereira, & Baker, 2020). The concept of such an abstract object space thus offers a framework for understanding the importance of linkage between the mind and the physical world (Love & Roads, 2021; Nosofsky, Sanders, Meagher, & Douglas, 2018; Riesenhuber, 2020). However, regarding the fundamental question of how the brain produces the mind (Shiffrin, Bassett, Kriegeskorte, & Tenenbaum, 2020), it is still unclear, in particular, how the abstract and complex multi-dimensional object space is physically implemented in a two-dimensional cortical surface of our brain (Bao, She, McGill, & Tsao, 2020; DiCarlo & Cox, 2007).

A candidate cortical region for the implementation of the object space is human ventral temporal cortex (VTC), a large area dedicated to the complex task of object recognition and containing numerous subregions with distinct selectivities for specific object categories (Grill-Spector & Malach, 2004; Grill-Spector & Weiner, 2014; Hasson, Levy, Behrmann, Hendler, & Malach, 2002; Op de Beeck, Haushofer, & Kanwisher, 2008). In this region, neurons specialized for processing specific categories of objects are clustered into discrete brain subregions, e.g., lateral occipital complex (LOC), fusiform face area (FFA), parahippocampal place area (PPA), and extrastriate body area (EBA), with apparent selectivity discontinuities at the boundaries of these regions (Downing, Jiang, Shuman, & Kanwisher, 2001; Epstein & Kanwisher, 1998; Haxby, Gobbini, Furey, Ishai, Schouten, & Pietrini, 2001; Kanwisher, McDermott, & Chun, 1997; Nasr, Liu, Devaney, Yue, Rajimehr, Ungerleider, & Tootell, 2011; Weiner & Grill-Spector, 2013). In addition, all these specialized functional regions are distributed and even partially overlapped along the VTC, exhibiting a certain kind of large-scale topographical organization. For instance, within the VTC, it is known that the middle fusiform sulcus (MFS) separates the real-world small-size, animate, and central maps from the large-size, inanimate, and peripheral maps, respectively. Here we asked whether the cortical map of the VTC is the physical implementation of the object space and, specifically, what principles might govern the layout of such a map.

Theoretically, the object selectivities in VTC might be organized in any spatial layout by any principle as long as it can faithfully transform the high-dimensional object space into a two-dimensional cortical map. However, the remarkable systematicity and consistency of the layout across individuals indicates that biological constraints must be the core of such an organization principle. Proposed by Ramon y Cajal and further developed by many others (Chklovskii & Koulakov, 2004; Koulakov & Chklovskii, 2001; Weigand, Sartori, & Cuntz, 2017), the wiring cost minimization principle, which minimizes the axonal and dendritic wiring cost associated with long-range connections, has been considered as a leading factor deriving maps in the brain. Here we incorporated this principle into a hybrid self-organizing map (SOM) model to explain the emergence of the topographical organization in the VTC.

There are two essential features of our computational model. First, unlike previous SOM models that usually take as input the stimulus space (or pixel space) to simulate the lower visual cortex (Bednar & Miikkulainen, 2003; Durbin & Mitchison, 1990; Durbin & Willshaw, 1987; Kohonen, 1989; Konkle, 2021; Linsker, 1988), our SOM model takes a high-dimensional object representation space as the input and outputs a tuned map (i.e., an artificial cortical surface of the VTC), in which nearby units in the map project to nearby points in the object space (Figure 1). The high-dimensional object representation space is obtained via a pre-trained deep convolutional neural network (DCNN), the AlexNet, because numerous studies have demonstrated a striking similarity in the response profile and functional hierarchy between the AlexNet and human ventral visual cortex (e.g., Cichy, Khosla, Pantazis, Torralba, & Oliva, 2016; Guclu & van Gerven, 2015; Khaligh-Razavi & Kriegeskorte, 2014; Liu, Zhen, & Liu, 2020; Wen, Shi, Zhang, Lu, Cao, & Liu, 2018; Yamins, Hong, Cadieu, Solomon, Seibert, & DiCarlo, 2014). Second and critically, the parameter of our SOM is biologically-constrained by the lateral wring span (i.e., lateral projection span of axon and dendrite) of neurons in the human temporal lobe to specifically simulate wiring optimization in human VTC.

**Figure 1.**
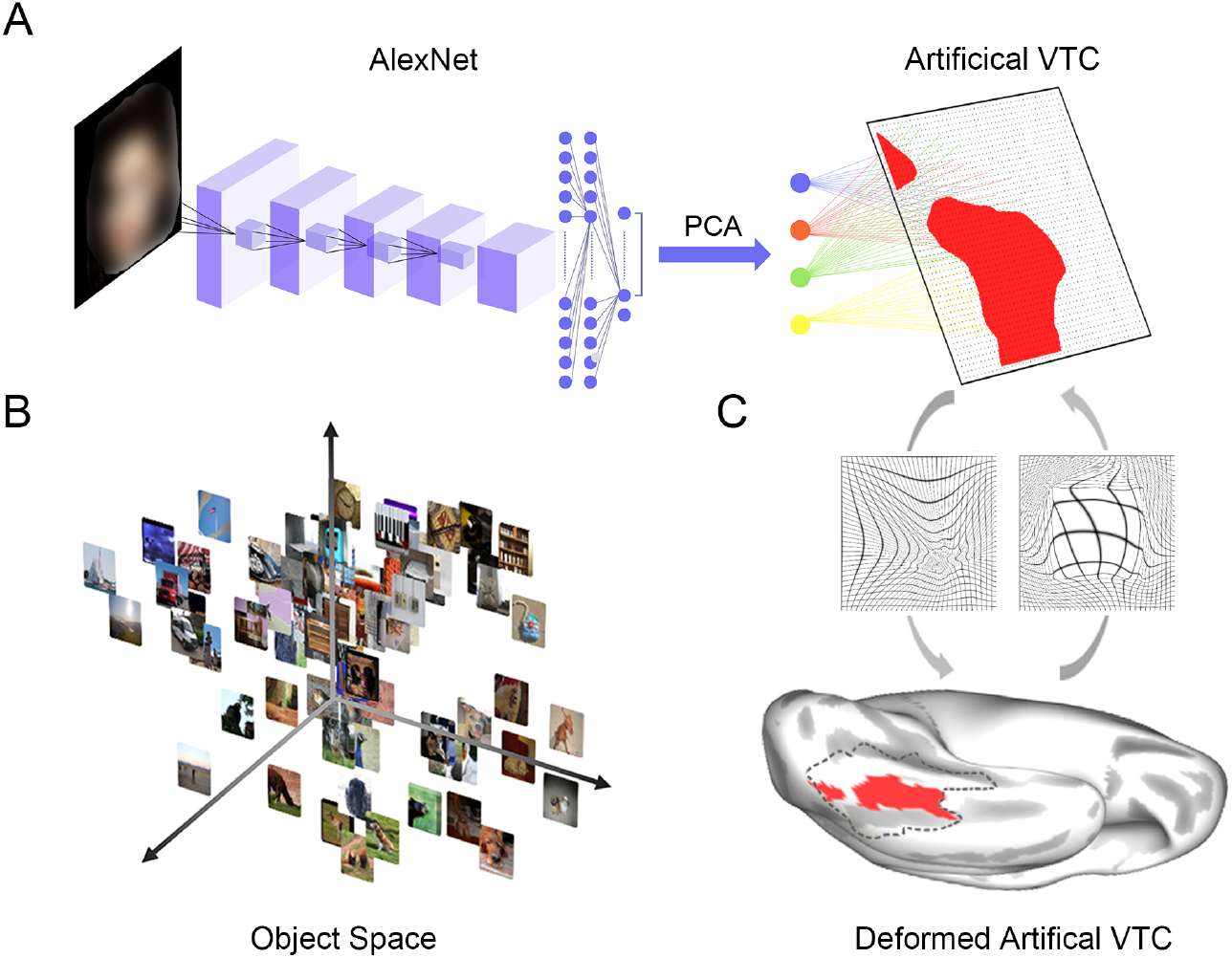
Model architecture and parameter selection. (A) Hybrid SOM model. The hybrid model combined a pre-trained AlexNet (left) with a SOM (right). The original FC3 layer of the AlexNet was reduced to a four-dimensional vector using the PCA, which then served as the input to the SOM (middle). Shown are an example input face image and the corresponding activation map in the SOM (i.e., face-selective regions). (B and C) The metaphorical high-dimensional object space derived from the pre-trained AlexNet (B) was physically implemented in the human VTC (C, bottom). The SOM learned a smooth 2-dimensional embedding (i.e., artificial VTCA) (A, right) to capture the structure of a 4-dimensional object space (A, middle), which simulated this physical implementation. To determine a biologically-constrained parameter sigma of the Gaussian function in the SOM, the 2-dimensional SOM (A, right) was co-registered to the flattened human VTC region in the HCP-MMP template (i.e., patch outlined by dark dashed lines) (C, bottom) using an SDR algorithm, and a deformation field (C, top left) was generated. The inverse of the deformation field (C, top right) also allowed for mapping a vertex in the human VTC onto the corresponding units in the SOM, through which an appropriate parameter sigma of the Gaussian function in the SOM can be biologically determined (see Methods for more details). Patch outlined by dark dashed lines represents the anatomically defined VTC region. Red regions within the patch represents the transformed face-selective regions in the SOM (for demonstration purpose only).

Having established the artificial VTC (hereafter referred to as the VTC-SOM), we then carried out two *in silico* experiments to systematically examine its functional specialization, topographic organization, and functionality reservation, including (1) qualitatively exploring the emergence of functional regions for four well-studied object categories (i.e., face, scene, object, and body) and their topological relations, and the similarity in the topographic organization of large-scale abstract functional maps (i.e., animacy, real-word object-size, and eccentricity bias maps) between the VTC-SOM and the human VTC identified with the functional magnetic resonance imaging (fMRI); and (2) quantitively testing of functionality reservation of the object-selective regions and networks in the VTC-SOM using the neurophysiological data recorded in the monkey inferior temporal cortex (ITC), a functionally homologous region to human VTC (Kriegeskorte, Mur, Ruff, Kiani, Bodurka, Esteky, Tanaka, & Bandettini, 2008). We found that the artificial VTC-SOM reproduced many topographic features of the human VTC and exhibited similar neural response patterns and functional hierarchy as those in monkey ITC. In short, our findings suggest that the wiring cost minimization principle in conjunction with the wiring span in the human temporal lobe is sufficient to determine the emergence of the map of object selectivity in the visual temporal lobe.

## Results

### Building an artificial VTC

#### SOM model architecture

We developed a hybrid computational model to simulate how the 2-dimensional cortical surface implements the high-dimensional object space. This hybrid computational model consisted of two modules (Figure 1A). The first module was a pretrained AlexNet (Figure 1A, left), which is used to obtain the high-dimensional object representation space (Figure 1B). Here, for computational simplification, the high-dimensional object space was reduced to a 4-dimensional object space using the Principal Component Analysis (PCA) and served as the input to the SOM (Figure 1A, middle). The second module was a SOM model equipped with the wiring cost minimization principle (Figure 1A, right). The SOM is not only a biologically plausible pattern-formation process (Obermayer, Blasdel, & Schulten, 1992), but also a simple algorithm enabling spatially adjacent units in a 2-dimensional lattice to project to nearby points in the object representation space according to the competitive-cooperative learning rule, which can minimize the overall wiring cost.

In brief, at each time step, a unit ***s*** at the location *r* in the lattice whose weight vector ***w***_*r*_ was closest to the pattern of a given input ***ξ*** randomly sampled from the object space, was thus defined as a winner unit (i.e., “winner-take-all” competition).

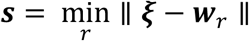

here, ***w***_*r*_ was a 1×4 vector of the connection weights between the unit in the 2-dimensional lattice and the four-dimensional feature vector representing the object space. And the neighbors of the winner unit ***s*** in the lattice were updated with this formula:

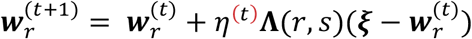

where **Λ**(*r, s*) was the neighborhood function, given by a Gaussian form:

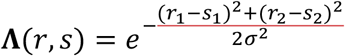

here, the parameter sigma (*σ*) is the full width at half maximum (FWHM) of the Gaussian function. As evident from the formula, **Λ**(*r, s*) was also a weighting function, and the parameter sigma determined the number of neighbors of the winner unit, so that only a few neighbor units in the lattice responded to each input. The closer a neighbor unit was to the winner unit, the more its weights got updated. The farther away a neighbor unit was from the winner unit, the less it learned. In this way, spatially neighboring units cooperated in the learning procedure to form a continuous cluster in the lattice (see Methods for more details).

#### Biological constraints

Next and importantly, a critical step in our model implementation was determining the value of parameter sigma. Inspired by the biological findings of the lateral activity spreads of the cortical neuron in local cortical circuits, **Λ**(*r, s*) can be treated as a rough approximation of the cortical point spread (CPS) function (McILWAIN, 1975; Park, Cha, & Lee, 2013), which was determined by the lateral projection span of neurons (Gilbert & Wiesel, 1983; Grinvald, Lieke, Frostig, & Hildesheim, 1994). Therefore, in our SOM model, we set the parameter sigma as the lateral wiring span of neurons in the human temporal lobe and kept it constant throughout the model training.

Then, based on the axonal data (Mohan et al., 2015) and dendritic data (Benavides-Piccione, Ballesteros-Yáñez, DeFelipe, & Yuste, 2002) of the human temporal lobe, we calculated a local lateral projection span of neurons in the human temporal lobe, which approximated a maximum length about 2.7 mm. We next registered the 2-dimensional lattice to the human VTC region in the Multi-Modal Parcellation (MMP) template using a Symmetric Diffeomorphic Registration (SDR) algorithm (Figure 1C) (Avants, Epstein, Grossman, & Gee, 2008), and estimated a parameter sigma (i.e., 6.2) matching the maximum wiring length of neurons in the human temporal lobe (see Methods for more details).

After the model training, the SOM learned a smooth 2-dimensional embedding to capture the structure of the object space, through which we obtained an artificial cortical surface of the human VTC (i.e., VTC-SOM).

### *In silico* Experiment 1: Spatial layout

#### Similarity in the fine-scale object-selective regions

After obtaining the VTC-SOM, we first examined the emergence of four classical object-selective regions (i.e., face-, scene-, object-, and body-selective regions) and their topological relations at the regional level. To do this, we fed the stimuli of four categories (face, place, body, and object) used for the fMRI study in the HCP dataset to our trained model, and obtained the activation map for each category in the VTC-SOM. We found that units selective for a particular category were clustered into patches (i.e., functional regions) (Figure 2A). Importantly, these functional regions showed a certain fine-scale topological arrangement. We then used the same SDR algorithm to transform the VTC-SOM to the left and right human VTC of the HCP-MMP template separately, and quantitively inspected the similarity in the topological organization of functional regions between the VTC-SOM and human VTC. Since the topology of the VTC regions did not differ greatly between the two hemispheres, Figure 2B only showed the comparison between VTC-SOM and human VTC in one hemisphere. Approximating to the functional organization of the human VTC identified with the HCP fMRI dataset, there were also two face-selective regions in the VTC-SOM (Figure 2B, upper left), and the two body-selective regions were located more lateral than the face-selective regions (Figure 2B, lower left). And the face- and place-selective regions were also adjacent to each other in the VTC-SOM (Figure 2B, lower right). Similarly, there were two object-selective regions in the VTC-SOM (Figure 2B, upper right). One resided in two body-selective regions, and the other was located adjacent to the place-selective regions. Though the place-selective regions seemed to be less laterally in the VTC-SOM than in the human VTC, the topological relations among functional regions were almost the same between VTC-SOM and human VTC. In sum, visual inspection suggests a similar spatial layout among functional regions between the VTC-SOM and human VTC (Figure 2B, central).

**Figure 2.**
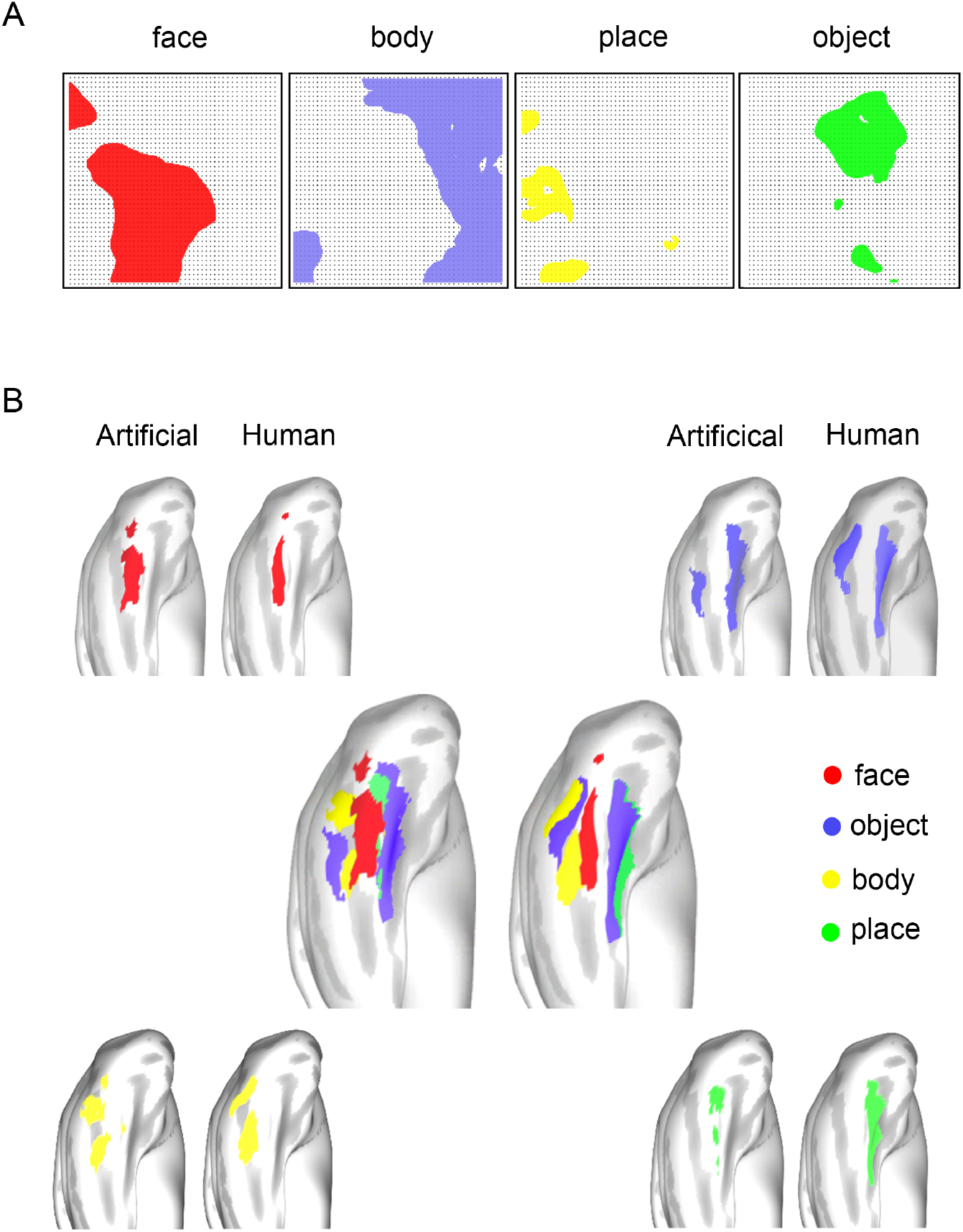
Similarity in the fine-scale object-selective regions between the VTC-SOM and human VTC. (A) Four object-selective regions in the VTC-SOM identified with the stimuli from the HCP dataset. Dark dots represent units in the SOM. Face-, body-, place-, and object-selective regions are shown from left to right, respectively. (B) These object-selective regions obtained in the VTC-SOM were transformed and mapped onto the left hemisphere of the human brain. The same object-selective regions identified with the human fMRI data were also displayed side-by-side for visual inspection. The spatial arrangements of the category-selective regions between the VTC-SOM and human VTC were similar (central). Face- (upper left), object- (upper right), body- (lower left), and place- (lower right) selective regions are colored by red, purple, yellow, and green, respectively.

#### Similarity in the large-scale abstract functional maps

One may argue that the emergence of the category-selective regions in the VTC-SOM was predictable because objects had already been classified into categories by the AlexNet within our computation model. However, this argument cannot explain the similarity in topological organization between the VTC-SOM and human VTC. Further, besides the fine-scale functional organization, human research and neurophysiological studies have also revealed large-scale functional maps in the VTC (for reviews, see Grill-Spector & Weiner, 2014; Op de Beeck, Haushofer, & Kanwisher, 2008). For instance, the animacy map, real-world object-size map, and eccentricity bias map are all aligned to the mid-fusiform sulcus (MFS) (patches outlined by dark lines in Figure 3), reflecting more abstract functional representations implemented in larger spatial scales across the VTC, which can generate broader categorical distinctions (Hasson, Levy, Behrmann, Hendler, & Malach, 2002; Haxby, Guntupalli, Connolly, Halchenko, Conroy, Gobbini, Hanke, & Ramadge, 2011; Konkle & Oliva, 2012). The coexistence of the fine-scale and large-scale functional organizations could increase the efficiency and flexibility of category processing in the VTC. If our computational framework is foundational in determining the topographic organization of the VTC, we should expect the emergence of large-scale abstract functional maps.

**Figure 3.**
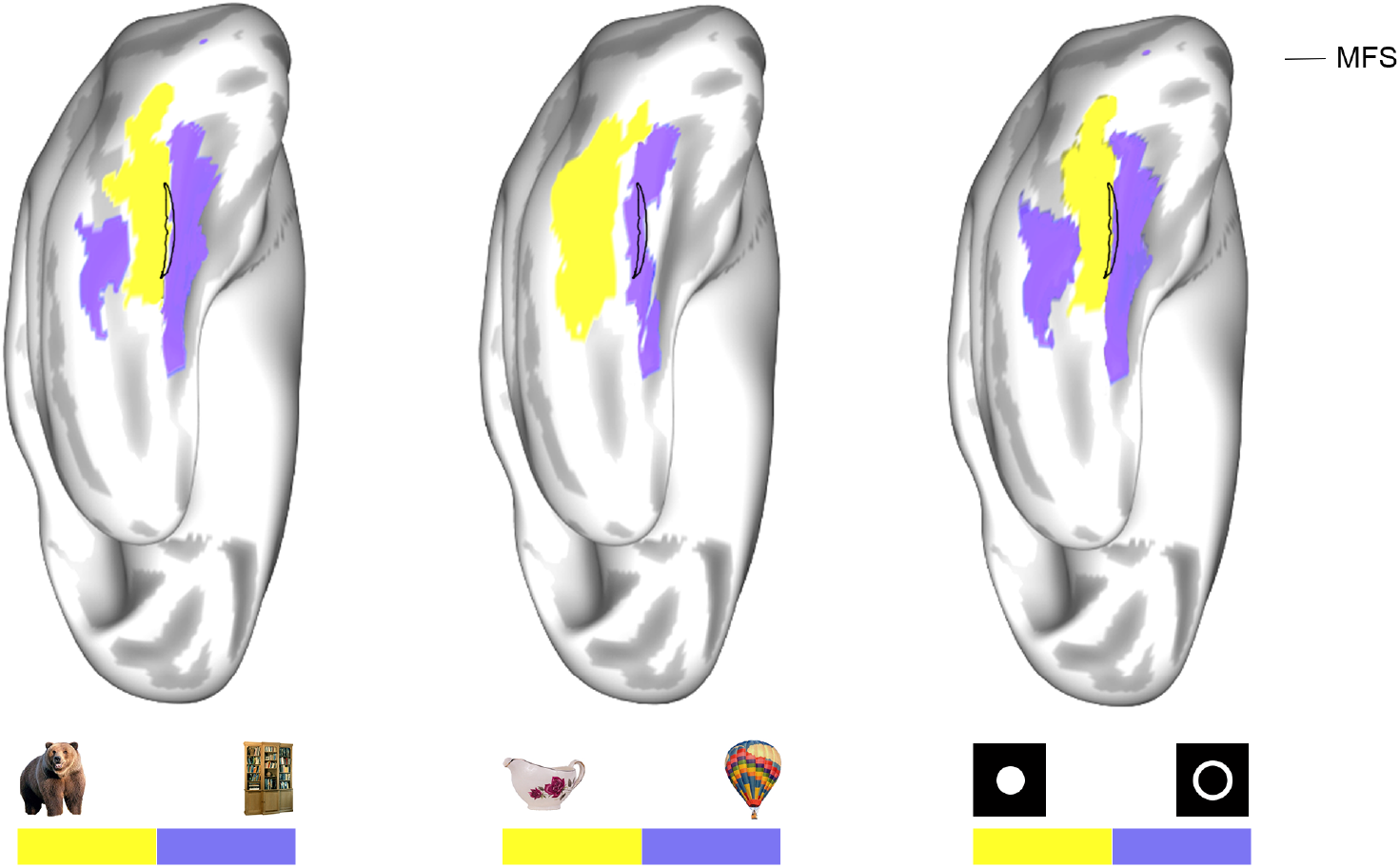
VTC-SOM reproduces large-scale abstract functional maps between the VTC-SOM and human VTC. The animacy (left), real-world object-size (middle), and eccentricity (right) maps in the VTC-SOM obtained using the animacy, real-word object-size, and the home-made eccentricity stimulus-sets, are transformed and mapped onto the left hemisphere of the human brain. Visual inspection suggests, the animate, small-size, and central maps in the VTC-SOM were also well separated from the inanimate, big-size, and peripheral maps, respectively, by the MFS. Patches outlined by dark lines represent the MFS. Animate, small-size, central maps are colored by yellow, whereas inanimate, big-size, peripheral maps are colored by purple.

To verify this prediction, we used images of the animacy and real-word object-size stimulus sets from the Konkle lab database (Konkle & Caramazza, 2013; Konkle & Oliva, 2012) as the input to our computational model and obtained the animacy map and real-world object-size map in the VTC-SOM. Similarly, we generated a set of ring stimuli with different eccentricities to obtain the eccentricity bias map. Again, we registered the three maps to the human VTC template using the same SDR algorithm. We found that these three maps were well separated by the MFS, like those found in human VTC (Figure 3). In short, the artificial VTC built by our computational model was similar to human VTC at the level of both fine-scale object-selective regions and large-scale abstract functional maps.

### *In silico* Experiment 2: Representation

Because functionality is determined mainly by anatomy, we expected that the VTC-SOM should represent objects in the same way as the VTC. To test this conjecture, we compared the functional representation of the VTC-SOM with that of monkey ITC at the neuronal level.

#### Similarity in functional representation

To examine the similarity in the representation of objects, the same images of real-world objects were presented to the VTC-SOM and monkey ITC. The stimuli contained 51 objects from six different categories, each presented at 24 different views. The neural responses to each stimulus were recorded from 482 neurons distributed in multiple patch networks of the monkey ITC, including face-, body-, stubby-, and spiky-selective patch networks (see Methods for more details). With the representational similarity analysis (RSA) (Kriegeskorte, Mur, & Bandettini, 2008), we carried out a Pearson correlation analysis on the response patterns between objects to obtain a representational similarity matrix (RSM) for the monkey ITC (Figure 4, right). Accordingly, we fed the same stimuli into our computational model and obtained an RSM for the VTC-SOM (Figure 4, left). Visual inspection of these two RSMs suggested a similar categorical representation in VTC-SOM and monkey ITC. Quantitative correlation analysis on the two RSMs revealed a substantial correlation (r = 0.63, *p* < 0.001), indicating the striking match in representation structure between the VTC-SOM and monkey ITC.

**Figure 4.**
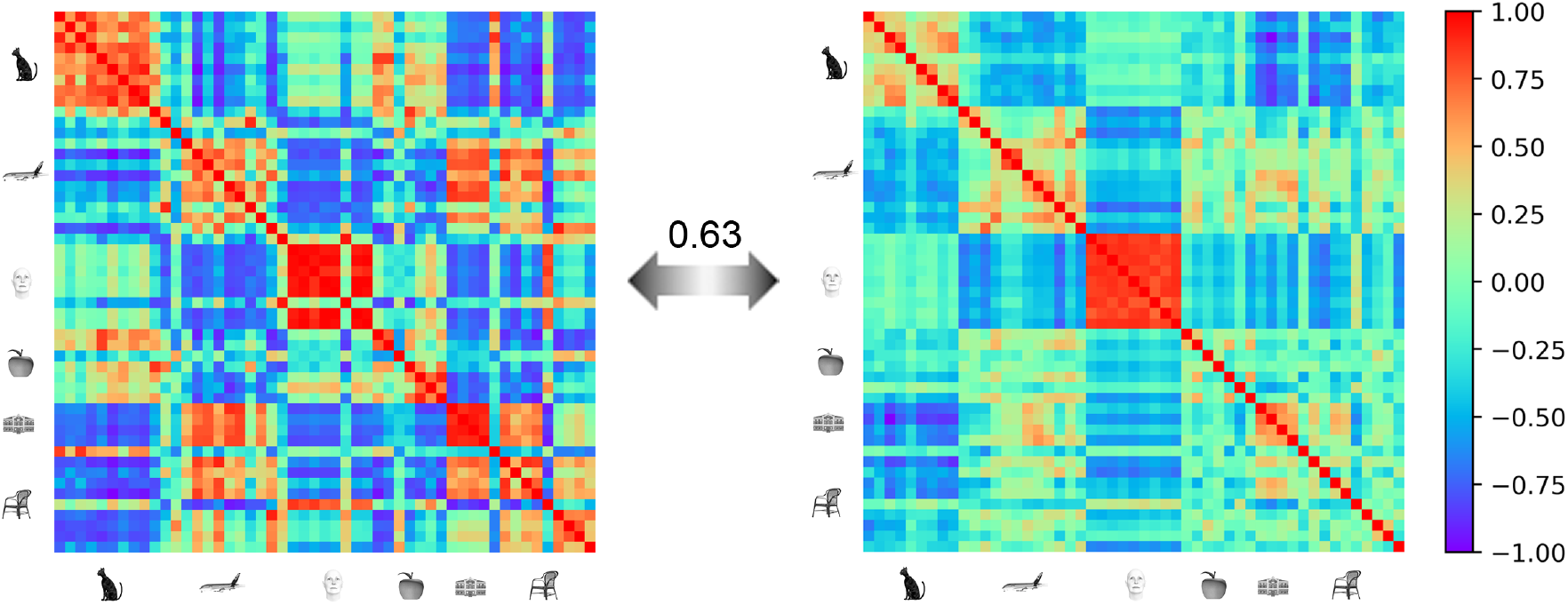
Similarity in the functional representation between VTC-SOM and monkey ITC. Model RSM for the VTC-SOM (left) and neural RSM for the monkey ITC (right).

#### Similarity in the functional representation within object-selective regions

Further, we examined the representation similarity within object-selective regions between the VTC-SOM and monkey ITC. To do this, we first identified ten most-preferred and ten non-preferred stimuli for each of the four patch networks in monkey ITC by computing the average responses of neurons in that patch network based on the single-unit data. Then, we fed these stimuli to our model and obtained their activation maps in the VTC-SOM. Units showing stronger activation for the preferred stimuli than non-preferred stimuli were defined as the face, body, spiky, and stubby network in the VTC-SOM separately, which corresponded to the patch networks in the monkey ITC accordingly. Finally, we conducted the RSA analysis to investigate whether the functional representation was similar within category-selective regions at the neuronal level between the VTC-SOM and monkey ITC. Indeed, we found strong correlations between VTC-SOM and monkey ITC RSMs in the face-, body-, spiky-, and stubby-selective regions (r = 0.76, *p* < 0.001; 0.77, *p* < 0.001; 0.53, *p* < 0.001; 0.74, *p* < 0.001, respectively) (Figure 5). These correlations again indicated that the functional representation of monkey ITC was preserved in VTC-SOM within object-category regions.

**Figure 5.**
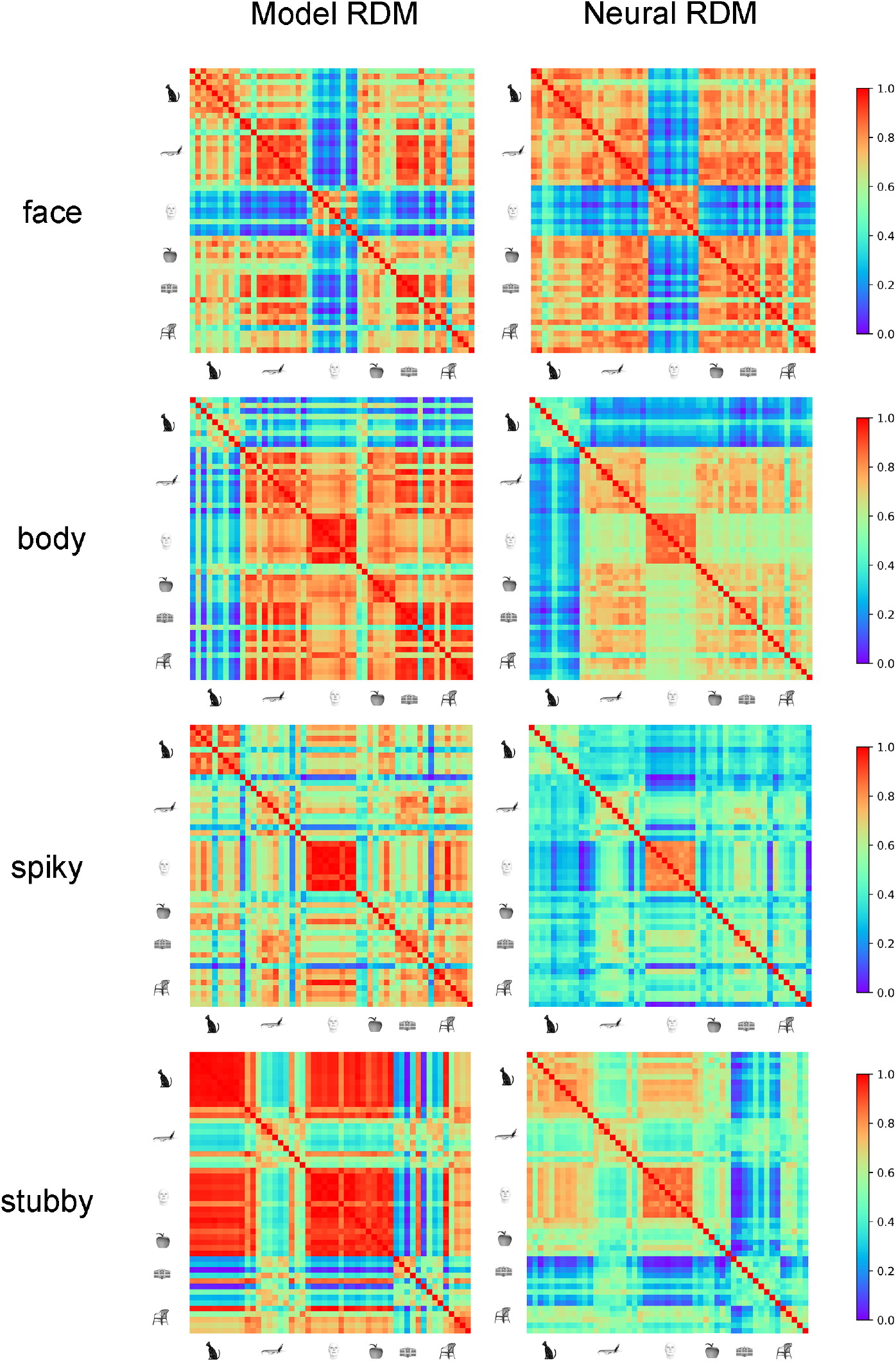
Similarity in the functional representation within object-selective regions between VTC-SOM and monkey ITC. Shown from top to bottom are model and neural RSMs for the face-, body-, spiky-, and stubby-selective regions in the VTC-SOM (left column) and monkey ITC (right column), respectively.

#### A hierarchy of increasingly view-invariant representations

Interestingly, multiple regions in the VTC-SOM were identified that showed selective responses to just one object category (Figure 2). Studies from neurophysiological studies suggest that in some patch networks of the monkey ITC, e.g., face patch network, neurons in posterior patches were view-specific and those in the most anterior patch view-invariant (Bao et al., 2020). We thus wondered whether such hierarchy in view invariance was also present in our VTC-SOM. To do this, we took two face stimuli, each of which contained 8 different views, as examples to explore the view-invariant representation within the three face-selective regions of the face network in the VTC-SOM. Specifically, for each of the three face-selective regions in the VTC-SOM, we fed the 8 views of each face stimulus into our model and computed an RSM from units’ activations to the views. As illustrated in Figure 6A, there were apparent differences in view invariance between three face-selective regions. One face-selective region, corresponding to the posterior face patch in monkey ITC, was in part view-specific to face stimuli. Another face-selective region, corresponding to the anterior face patch in the monkey ITC, was nearly view-invariant to face stimuli. This hierarchy of increasingly view-invariant representations was apparently face-specific, because for the non-face stimuli (e.g., airplane and vase), no such trend was found in these regions (Figure 6B). In short, by having multiple regions for the same object category, our VTC-SOM also achieved both view-dependent and view-independent representational abilities, similar to monkey ITC.

**Figure 6.**
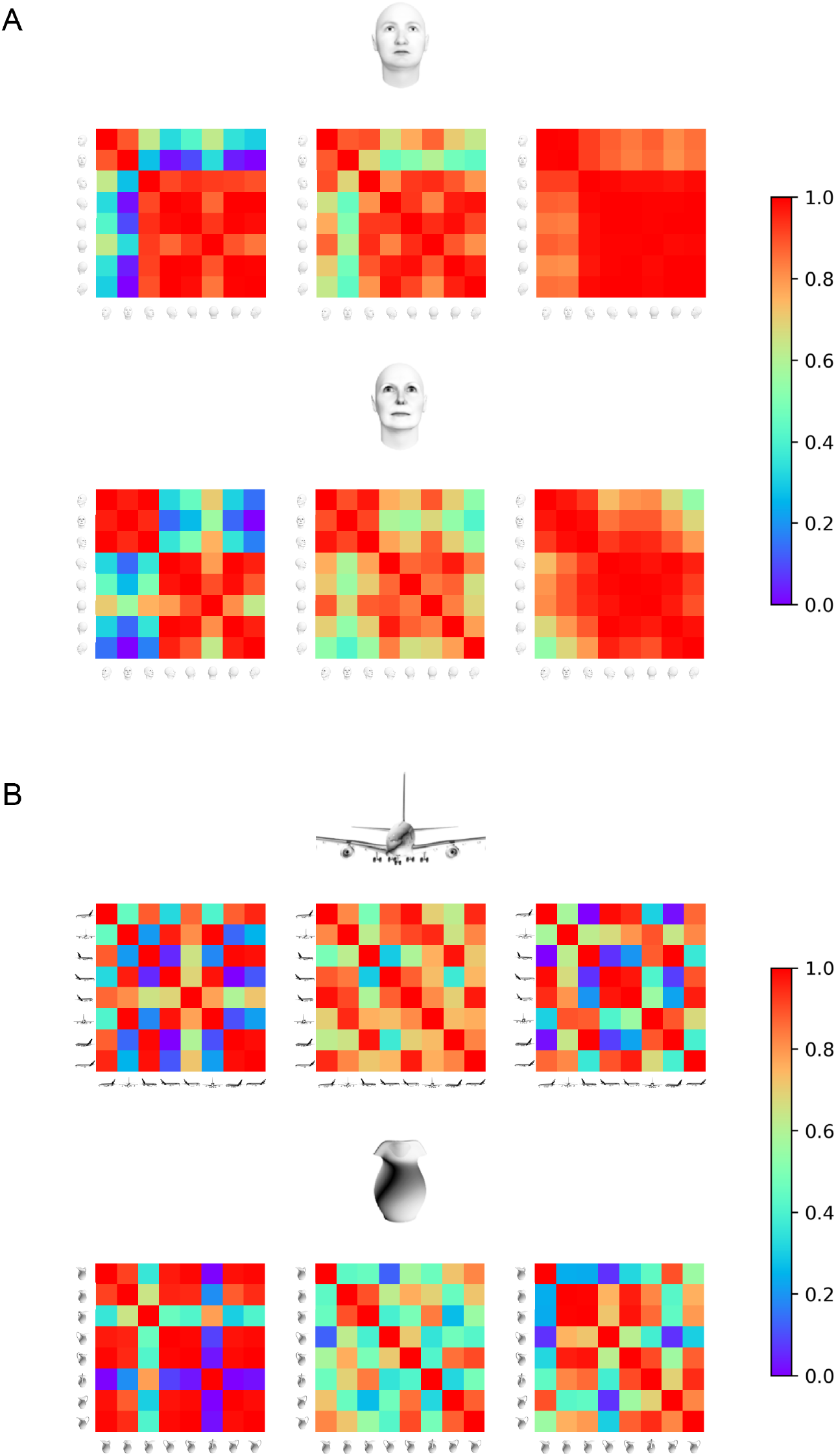
A hierarchy of increasingly view-invariant representations in the face network of the VTC-SOM. (A) RSMs for the face stimuli within the three face-selective regions of the face network in the VTC-SOM. There was an apparent increase in the view-invariance between three face-selective regions. For both face stimuli, there were significant differences in the view-invariance between the first (left-most panels) and third (right-most panels) face-selective regions (KS-test, all *ps* < 0.05). (B) RSMs for the non-face stimuli within the three face-selective regions of the face network in the VTC-SOM. No significant increase in the view-invariance was found between three face-selective regions (KS-test, all *ps* > 0.05).

### The critical role of the wiring span

According to the wiring optimization principle, maps develop to minimize wiring costs associated with long-range connections (Chklovskii & Koulakov, 2004); therefore, maps can arise in any cortices of the brain, regardless of the nature of its content or the complexity of its computations. However, the specific layout of the VTC is drastically different from those in early sensory cortices, for example, suggesting that the wiring span, the only parameter in the principle, may play a critical role in shaping the spatial layout of the VTC. To test this intuition, here we varied the parameter sigma in a broad range and examined whether the resulting VTC-SOMs showed different functional organizations. In our model, the object space constructed by the first four principal components was projected to a 2-dimensional lattice. For better visualization, here we only illustrated the projection of an object space constructed by the first two principal components (i.e., the PC1-PC2 space).

For each unit at the position ***r*** in the lattice, the arctangent value of the ratio between the first two weights of its ***w***_*r*_ corresponded to the polar angle *θ*_*r*_ in the PC1-PC2 space, given by

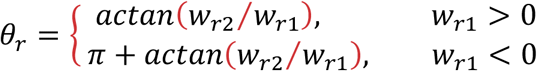

Besides the sigma of 6.2 used in the model, here we also trained the SOM with other values: 0.1, 0.7, 1.7, 3.1, 9.3, and 10.0, while keeping the input and the lattice architecture unchanged. In the SOM with different parameter sigma, we calculated every unit’s polar angle. As shown in Figure 7, a transition in the functional organization can be observed as a function of the sigma values. When the sigma value was small, the units in the SOM showed an unstructured salt-and-pepper organization. As the sigma value increased, a structured pinwheels-like pattern began to emerge and gradually changed to a pattern with many pinwheels, and finally changed to a pattern with only one big pinwheel. That is, only when the sigma value matched the wiring span of neurons in the human temporal lobe, the VTC-SOM could exhibit similar functional organization to that of the human VTC. Therefore, it is the wiring span that determines the functional organization of the VTC as it is.

**Figure 7.**
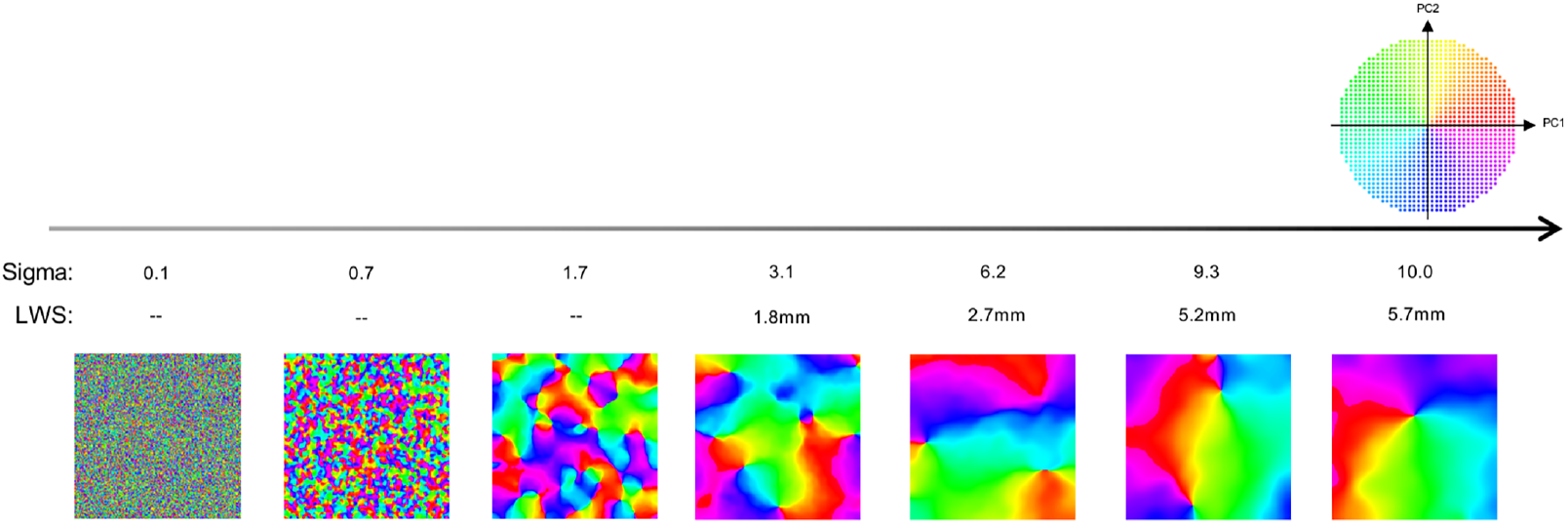
Transition in the functional organization of the VTC-SOM as a function of the parameter sigma. The functional organization of the VTC-SOM for different sigma values. As the sigma value increased, the functional organization in the VTC-SOM changed from an unstructured salt-and-pepper organization (left) to a structured pinwheels-like arrangement and gradually changed to a pattern with many pinwheels (middle), and finally changed to a pattern with only one big pinwheel (right). For better visualization, all units in the SOM were colored by their polar angles in the PC1-PC2 space. The color scheme was shown in the upper right inset. Different colors indicate different polar angles in the PC1-PC2 space. LWS denotes the lateral wiring span. “--” indicates that the LWS value cannot be computed when the estimated number of vertices, corresponding to the sigma value of 0.1, 0.7, or 1.7, is less than one vertex.

## Discussion

According to Marr’s tri-level hypothesis (Marr, 1982), to achieve the computational goal of rapidly recognizing miscellaneous objects in the environment despite large variations in their appearance, our mind, at the algorithm level, develops an algorithm of representing objects in a high-dimensional object space to provide separable category information and enable the extraction of different kinds of information necessary for various levels of the visual processing. At the implementational level, mounting evidence from human fMRI and monkey neurophysiological studies has revealed that the VTC serves as the physical substrate to implement the abstract and complex object representation space (DiCarlo, Zoccolan, & Rust, 2012; Felleman & Van Essen, 1991; Hegdé & Van Essen, 2007; Kobatake & Tanaka, 1994; Livingstone & Hubel, 1988; Logothetis & Sheinberg, 1996; Mishkin, Ungerleider, & Macko, 1983), and develops different functional regions selectively responsible to specific object categories, such as faces and places, with a certain kind of large-scale topographic organization (Grill-Spector & Weiner, 2014; Haxby, Gobbini, Furey, Ishai, Schouten, & Pietrini, 2001; Kanwisher, McDermott, & Chun, 1997; Nasr, Liu, Devaney, Yue, Rajimehr, Ungerleider, & Tootell, 2011; Weiner & Grill-Spector, 2013). Here, we propose a computational framework to explain how the abstract and complex object representation space at the algorithmic level is faithfully implemented in a biological system both anatomically and functionally. Inspired by the principle of wiring cost minimization, we built a hybrid SOM model with biological constraints on the model parameter and simulated an artificial cortical surface of the human VTC (VTC-SOM). In two *in silico* experiments with the empirical brain imaging and single-unit data, our VTC-SOM can predict the emergence of several fine-scale functional regions (including the classical object-selective regions, as well as the newly-found spiky and stubby regions) and the boundary (i.e., MFS) that indicates transitions in large-scale abstract functional maps (animate vs. inanimate, real-word large-size vs. small-size, central vs. peripheral), with no significant loss in functionality (i.e., categorical selectivity, a hierarchy of increasingly view-invariant representations). In sum, our study bridges the important gap between the algorithm and implementation levels for object recognition by providing a computationally simple and biologically realistic model for the emergence of the topographic organization of object selectivity in the VTC.

In a biological brain, along the hierarchy of the visual stream, the wiring cost minimization is a general principle that explains a strong tendency of cortices to exhibit local clustering (Chklovskii & Koulakov, 2004; Koulakov & Chklovskii, 2001), because the minimization of wire length will ensure that neurons representing similar features lie close together and thus form functional modules and maps in the cortex. These functional modules and maps can be understood as a result of dimension-reduction mapping from high-dimensional representation space to the cortical surface. Theoretically, there are many dimension-reduction methods to achieve this goal. Some dimension-reduction methods such as PCA, t-Distributed Stochastic Neighbor Embedding (tSNE), and Multiple Dimensional Scaling (MDS) map the multi-dimensional input space into 2-dimensional coordinates without considering their physical implementations, and thus for which they still simulate cognitive processes within the algorithm level. In other words, there is no transition from the algorithm level to the implementation level with these methods.

In contrast, the SOM model operates in an opposite way by mapping a 2-dimensional lattice into the input space (Durbin & Mitchison, 1990; Konkle, 2021), while preserving the most important topological and metric relationships of objects in the high-dimensional object space. The SOM model assumes a regularly, locally connected cortical surface and learns an embedding in the input space, which makes it well-suited to answer the question of how a 2-dimensional cortical surface enables the fundamental ability to represent a multi-dimensional object representation space in our study. When a 2-dimensional lattice is projected into a multi-dimensional object space, nearby units in the map are projected to nearby points in the object space; therefore, the SOM model is a biologically plausible model that can achieve local wiring cost minimization (Durbin & Willshaw, 1987; Kohonen, 1989). Though the SOM models have been successfully applied to predict the appearance of ocular dominance, orientation preference, and direction preference maps in the early visual cortex (Bednar & Miikkulainen, 2003; Durbin & Mitchison, 1990; Konkle, 2021; Linsker, 1988), our study is the first attempt to use the SOM to model the functional organization of higher visual cortex, the VTC, both anatomically and functionally.

Another merit of our SOM model is that we take the lateral wiring span as the critical biological constraint on the model parameter. This is inspired by another biological findings of cortical point spread function, in which neural activities triggered by external stimuli spread laterally from the initial firing neuron to surrounding neurons, through its axonal projections to the nearby neurons (McILWAIN, 1975; Park, Cha, & Lee, 2013). The spatial extent of this lateral projection thus serves as an important biological constraint for the cooperative process (i.e., parameter sigma of the Gaussian function in the model) of the SOM (Gilbert & Wiesel, 1983; Grinvald, Lieke, Frostig, & Hildesheim, 1994). In fact, adjusting the sigma value of the Gaussian function in our SOM model can lead to different spatial patterns, indicating that the lateral wiring span is a critical factor in determining the functional organization of the human brain. Further, only a moderate value of parameter sigma that matches the lateral wiring span in the human temporal lobe can result in a functional organization similar to the human VTC. Intuitively, it seems to be inconsistent with the wiring cost minimization principle because the sigma value in our SOM has a moderate size rather than being minimal. However, from the perspective of computational neuroscience, advanced information-processing capabilities are usually associated with the greater-than-minimal wiring cost, which reflects a trade-off between network cost and efficiency in computation and could enable a system simultaneously accommodating generality and specificity. In line with this perspective, it has been found that representations of object categories in the VTC show both generality and specificity properties. Responses in the VTC generalize across exemplars within the same category and tolerate transformations of their low-level features such as position and size, while distinguishing between categories with similar physical attributes and configurations. Therefore, unlike the previous study (Lee, Margalit, Jozwik, Cohen, Kanwisher, Yamins, & DiCarlo, 2020), our model demonstrates that the functional organization in the VTC is determined by the proper wiring span in the human temporal lobe, which might be optimized by evolutionary and/or developmental needs to accommodate both the generality and specificity.

One beauty of our model is that it does not need to make any assumptions about the functional and architectural differences along the hierarchy of the human VTC. For instance, one puzzling feature of the VTC is that some object-selective regions (e.g., faces, places, and general objects) come in pairs, with one located on the ventral temporal cortical area and the other on the lateral occipital cortical area (Beauchamp, Lee, Haxby, & Martin, 2002; Hasson, Avidan, Deouell, Bentin, & Malach, 2003; Levy, Hasson, Avidan, Hendler, & Malach, 2001). Based on the dual pathway model (Mishkin, Ungerleider, & Macko, 1983), an organizational principle is proposed that regions on the lateral area are more sensitive to location information and regions on the ventral area to shape information (Beauchamp, Lee, Haxby, & Martin, 2002; Hasson, Avidan, Deouell, Bentin, & Malach, 2003). Our VTC-SOM did not make such an assumption, yet the feature of paired object-selective regions was also present. Therefore, this feature likely derives from the nature of the object space, rather than predisposed architectural and functional differences in the VTC. Similarly, our model does not support organizational principles of the temporal association of objects based on everyday experiences (e.g., co-occurrence of the table and chair) (Erickson & Desimone, 1999; Messinger, Squire, Zola, & Albright, 2001; Sakai & Miyashita, 1991), the computation required to recognize particular object categories (e.g., fine-grained visual acuity for recognizing faces) (Levy, Hasson, Avidan, Hendler, & Malach, 2001; Malach, Levy, & Hasson, 2002), or the conceptual knowledge about objects (e.g., animacy) (Warrington & McCarthy, 1983), simply because these factors are absent when training the AlextNet to construct the object space and then mapping it to a 2-dimensional lattice. In short, our model suggests that the principle governing the topological organization of object selectivity in the VTC might be much simpler than originally hypothesized.

Finally, unlike a biological brain, our model is computationally transparent and entirely accessible to visualization and manipulation, which enables conducting various *in silico* cognitive neuroscience experiments. For instance, we will be able to formulate hypotheses about the functional organization at multiple spatial scales in the brain across species and across cortices that can be validated by observing the effects on predicted maps resulting from changes to the model parameter. We could also address fundamental questions about brain computations in the developing brain by manipulating critical model parameters to simulate the entire process of development. In addition, lesioning the model components could unravel features critical for the formation of brain functions. Even more promisingly, our model offers a new tool for researchers to explore how other multi-dimensional abstract representation space, e.g., knowledge representation space, value space, or decision space for action planning, is implemented in the two-dimensional cortical surface of the brain.

## Acknowledgments

This work was supported by the National Natural Science Foundation of China (31861143039), and National Key R&D Program of China (2019YFA0709503).

## Author Contributions

Y.Z. and J.L. conceived the study design. Y.Z. built computational models and performed analyses. P.B. provided the single-unit data. K.Z. and J.L. provided technical advice on model implementation. K.Z., Y.Z., and J.L. interpreted the data. K.Z., Y.Z., and J.L. wrote the paper.

## Data and code availability

Data and custom code are available from the corresponding author upon reasonable request.

## Methods

### Building an artificial VTC

#### Overall model architecture

The hybrid model consisted of two modules. The first module was a pretrained AlexNet, which was used to obtain a high-dimensional object representation space (Figure 1A, left). The second module was a SOM (Kohonen, 1989) with the wiring length of the human VTC as the critical biological constraint on the model parameter (Figure 1A, right).

#### Constructing a high-dimensional object space using the AlexNet

The AlexNet consisted of five convolutional and three fully connected layers (FC1– FC3). The FC3 was followed by a sublayer using a SoftMax function to output a 1×1000 vector, representing the probability of the visual input containing the corresponding object category (Krizhevsky, 2014).

First, by extracting a 1×1000 vector of the FC3 layer for each of the 50000 images of the training stimulus-set mentioned in 1.1, we constituted a 50000×1000 matrix. Then, for computational simplification, a principal component analysis (PCA) was conducted on the Z-scored matrix to reduce the dimensionality of the object space. An obvious inflection point of explanatory value was found at principal component 4. We thus retained only the first 4 PCs (Figure 1A, middle), which captured 47% of the response variance, to represent a four-dimensional object space, **Ξ**.

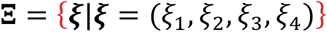

#### SOM model

In the SOM, each unit corresponded to a small piece of cortex containing many neurons and was labeled by its position in the lattice: ***r*** = (*r*_1_, *r*_2_). The weight vector ***w***_*r*_ characterized the property of the unit’s receptive field at position ***r***. The input features (1×4 vector) and all units in this lattice were fully connected. Therefore, the ***w***_*r*_ was also a four−dimensional vector whose components were ***w***_*rk*_ where *r* was the position of the unit and *k* was the index of the input features:

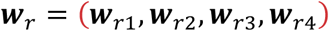

The pattern formation process determined by the SOM algorithm was discrete in both time and space. In each step of the training procedure, an element *ξ* was extracted randomly from the object space **Ξ** without repetition, and normalized by *ξ*⁄∥ *ξ* ∥. Then, the unit ***s*** = (*s*_1_, *s*_2)_) whose weight matrix ***w***_*s*_ of ***s*** was the closest to the input pattern *ξ*, was defined as the winner unit:

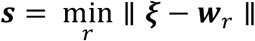

The weight vector ***w***_*r*_ was then updated by a dynamical equation, being constrained with a cortical point spread function determined by the lateral projection span:

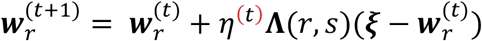

where *ŋ*^(*t*)^ is the learning rate, given by

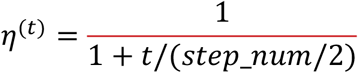

**Λ**(*r, s*) was the neighborhood function, characterized in a Gaussian form:

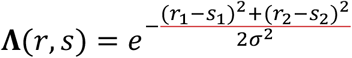

Here, the parameter sigma (*σ*) is the full width at half maximum (FWHM) of the Gaussian function. **Λ**(*r, s*) was also a weighting function, and the parameter sigma determined the number of neighbors of the winner unit ***s***, so that only a few neighbor units in the lattice responded to each input. It thus resulted in that the learned weights of the unit at the positions ***r***, which was adjacent to the winner unit ***s***, were close to the weights of the winner unit. The similarity in weights between units decreased with the increasing distance between ***r*** and ***s***. The parameter step_num was the total number of steps of the training iterations. At the end of each training step, the weight was normalized by

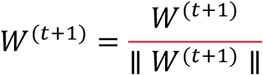

#### Defining the anatomical VTC template of the human brain

According to Grill-Spector and Weiner (2014), the lateral, posterior, medial, and anterior boundaries of the VTC were defined by the occipitotemporal sulcus (OTS), posterior transverse collateral sulcus (ptCoS), parahippocampal gyrus (PHG), and the anterior tip of the mid-fusiform sulcus (MFS), respectively. Following their delineation method, in the HCP-MMP template (Glasser et al., 2016), we combined eight anatomical areas (i.e., V8, FFC, PIT, PHA3, TE2p, PH, VMV3, and VVC) into a coherent region to serve as the human VTC template (Figure 1C, patch outlined by dark dashed lines).

#### Registration between the SOM and human VTC template

We used a Symmetric Diffeomorphic Registration (SDR) algorithm to establish the correspondence between units of the SOM and vertices of the human VTC template. The SDR method allowed for the accurate alignment of two maps when more significant deformations were needed, and the obtained deformation field was continuous. The SDR method also produced an inverse deformation field (Avants, Epstein, Grossman, & Gee, 2008).

Specifically, we first obtained the coordinates of all vertices in the VTC template and generated a smooth map by interpolation between the coordinates with the same values. We used the binary images of the SOM and VTC template for the alignment and chose the sum of squared differences as the loss function for the SDR method, through which we got the forward (Figure 1C, upper left) and inverse (Figure 1C, upper right) deformation fields between the SOM and VTC template and established the correspondence between them.

#### Determining a biologically-constrained parameter sigma for the SOM

Inspired by the biological findings of the lateral activity spreads of the cortical neuron in local cortical circuits, **Λ**(*r, s*) can be treated as a rough approximation of the cortical point spread (CPS) function (McILWAIN, 1975; Park, Cha, & Lee, 2013), which was determined by the lateral projection span of neurons (Gilbert & Wiesel, 1983; Grinvald, Lieke, Frostig, & Hildesheim, 1994). Therefore, in our SOM model, we set the parameter sigma as the lateral wiring span of neurons in the human temporal lobe and kept it constant throughout the model training.

First, based on the axonal data (Mohan et al., 2015) and dendritic data (Benavides-Piccione, Ballesteros-Yáñez, DeFelipe, & Yuste, 2002) in the human temporal lobe, the local lateral projection span of neurons in the human VTC approximated a maximum length about 2.7 mm. Then, we calculated a geodesic distance map (GDM) with a geodesic distance of 2.7 mm around each vertex. Separately for the left and right human VTC, we mapped the GDM of each vertex onto the SOM using the deformation field generated above, and computed the average number of units in the SOM that the GDM of one vertex occupied. Since the different alignment angles would lead to different results, we calculated the number of units in the SOM for each of the five alignment angles (60, 120, 180, 240, and 300 degrees). Next, we computed an average number of units occupied by one vertex. As a result, the average number of units in the SOM occupied by a vertex in the left and right human VTC were 532.6 and 570.3, respectively. When the parameter sigma was 6.2, most part of the distribution of the Gaussian function (i.e., height value greater than 0.1) occupied 553 units, which was between 532.6 and 570.3. Therefore, the parameter sigma in our SOM implementation was fixed to 6.2, which approximated the lateral projection span (2.7 mm) of neurons in the human temporal lobe.

#### Model training

Before the model training, the weights of the model were randomly initialized. The size of the SOM was set to 200×200 units. The parameter sigma of the Gaussian function was set to 6.2 and kept constant, reflecting the biological constraint of the lateral wiring span of neurons in the human temporal lobe. We used the natural images from the validation dataset of the ImageNet Large Scale Visual Recognition Challenge (ILSVRC) 2012 competition (50 images for each of the 1000 categories) to train the SOM. In each training step, the SOM learned the representation of an image in the four-dimensional object space *ξ*. The training process was iterated for a total of 200000 steps. At the end of the training, we obtained an artificial cortical surface of the human VTC (i.e., VTC-SOM).

### *In silico* Experiment 1: Spatial layout

#### Stimulus-set for identifying object-selective regions in the VTC-SOM and human VTC

To define the object-selective regions in VTC-SOM and human VTC, we used the stimuli of a working memory task from the HCP, which was combined with the category-specific representation task that can localize the object-selective regions. During the fMRI scanning, participants were presented with blocks of trials that consisted of pictures of places, tools (i.e., general objects), faces, and body parts (non-mutilated parts of bodies with no “nudity”), 61, 110, 111 and 78 pictures for each category, respectively (Van Essen, Smith, Barch, Behrens, Yacoub, Ugurbil, & Consortium, 2013).

#### Stimulus-sets for defining abstract functional maps in the VTC-SOM

To obtain the animacy map for the VTC-SOM, we used 120 animate images and 120 inanimate images from the animacy stimulus-set of the Konkle lab database (https://konklab.fas.harvard.edu/ImageSets/). Due to the lack of human images in the Konkle lab database, 40 images containing human information from the Algonauts Project Challenge 2019 dataset were added to the animate category.

To obtain the real-world object-size map for the VTC-SOM, we used 400 images from the real-word object-size stimulus-set of the Konkle lab database, 200 for each of the big- and small-size categories.

We also generated an eccentricity stimulus-set to obtain the eccentricity bias map. There were 40 eccentricity images (224 × 224 pixels), 20 for each of the central and peripheral categories. For the central-category stimuli, one solid disk was presented at the center of the image, whose radius ranged from 10 to 30 pixels. For the peripheral-category stimuli, one ring was presented at the center of the image. The outer radius ranged from 60 to 80 pixels, and the inner radius was fixed to 50 pixels.

#### *Human* f*MRI data for identifying object-selective regions in the human VTC*

The group-average fMRI data of the working memory task from the HCP were used to define the face-, place-, body-, and object-selective regions in the human VTC. There were two runs for the working memory task. Each run consisted of 8 task blocks (10 trials of 2.5 seconds each, for 25 seconds) and four fixed blocks (15 seconds). The four different stimulus types (i.e., place, tool, face, and body part) were presented in separate task blocks. The stimulus was presented for 2 seconds in each trial, followed by an inter-stimulus interval of 500 ms.

#### Identifying fine-scale object-selective regions in the human VTC

The HCP provided a group-average activation map (Cohen’s d map) for each of the four categories of the working memory task. For each category, vertices in the VTC with the Cohen’s d greater than 0.5 were defined as the category-selective vertices. In this way, we identified four object-selective regions in the human VTC that responded to face, place, body, and object stimuli, respectively (Figure 2B).

#### Obtaining fine-scale object-selective regions in the VTC-SOM

To define the object-selective regions in the VTC-SOM, we fed the same stimuli of the HCP working memory task into our trained model. Then for each stimulus, we obtained an activation matrix *A* of the VTC-SOM. Each element in *A* was represented as

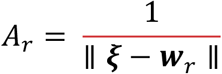

Then for each category, using a similar analysis as that used in the HCP fMRI dataset, we calculated a Cohen’s d map for this category, and the average Cohen’s d map for the other three categories. Subsequently, units that had the Cohen’s d greater than 0.5 and survived the independent T-Test (Bonferroni corrected p < 0.05) were defined as the units selective for this category. In this way, we obtained the face-, place-, body-, and object-selective regions in the VTC-SOM (Figure 2A).

Next, we aligned the human VTC template and VTC-SOM at multiple angles using the same SDR method and generated the forward and inverse deformation fields for each angle. We applied these deformation fields to the four object-selective regions in the VTC-SOM to obtain the transformed object-selective regions in the human VTC template (Figure 2B). As mentioned before, different alignment angles would lead to different results. We thus chose the angle, with which the transformed face-selective regions were best aligned to the face-selective regions identified with the HCP fMRI dataset in the same VTC template, as the final alignment angle for the current study. Importantly, this alignment angle and its corresponding deformation fields were used to transform other functional regions and maps throughout the following analysis.

Finally, we visually inspected whether the spatial locations and arrangement of the other three transformed object-selective regions in the human VTC template were similar to those identified with the HCP fMRI dataset.

#### Identifying large-scale abstract functional maps in the VTC-SOM

Using the same analysis described above, we obtained the animacy and real-world object-size maps in the VTC-SOM by feeding the images of the animacy and real-word object-size stimulus-sets to our trained model. Then, the simulated animacy and real-world object-size maps in the VTC-SOM were transformed onto the human VTC template using the same deformation field generated above. The eccentricity bias map of the VTC-SOM was obtained in the same way except that the custom eccentricity stimulus-set were used as the input to the pre-trained hybrid model. The similarity in the functional organization of these large-scale abstract functional maps between the VTC-SOM and human VTC was visually inspected (Figure 3).

## *In silico* Experiment 2: Representation

### Stimulus-set for single-unit recording and defining object-selective network in the VTC-SOM

The stimulus-set used for single-unit recording in the monkey ITC contained 51 stimuli from six different categories, including the animal, vehicle, face, vegetable, house, and object. Each stimulus was presented at 24 different views. The face stimuli were generated by the FaceGen software (https://www.facegen.com) using random parameters, and the stimuli for other categories were downloaded from https://www.3d66.com. The images at 24 views for each stimulus were generated using the 3DMAX software.

### Monkey single-unit data for testing the functionality of the VTC-SOM

Single-unit data were obtained from Bao et al. (2020), which were recorded in the ITC of a male macaque monkey using the stimulus-set mentioned above. During the experiment, the monkey was head-fixed and passively viewed the screen. Picture stimuli were presented on a CRT monitor in a random order. The screen size was 27.7° × 36.9°, the stimulus size was 5.7°, and the size of the fixation point was 0.2° in diameter. The position of the eye was monitored using an infrared eye-tracking system (ISCAN). Electrical signals were recorded from 483 neurons in the ITC, of which 10, 190, 67, and 216 neurons were from the face-, body-, stubby-, and spiky-selective patches, respectively. For more details on this study and data, see Bao et al. (2020).

### Testing the similarity in functional representation between VTC-SOM and monkey ITC

First, we calculated a neural RSM for the monkey ITC using the monkey single-unit data. The response value of each neuron for each image was defined as the baseline-subtracted responses in a window of 60-220 ms after image onset. The responses of each neuron to each stimulus were averaged across 24 views. Each element in the neural RSM represented the Pearson correlation between response patterns across neurons elicited by two stimuli.

Then, we fed the same images of the stimulus-set for single-unit recording to our trained model, and recorded the activations of the model units of the VTC-SOM. We also calculated an RSM for the VTC-SOM. Each element in the VTC-SOM RSM represented the correlation coefficient between activation patterns of units elicited by two stimuli.

Finally, we computed the Pearson correlation of the upper triangle between the VTC-SOM RSM and neural RSM of the monkey ITC to examine their similarity in functional representation (Figure 4).

### Testing the similarity in functional representation within object-selective regions

We used the same stimulus-set for single-unit recording to obtain the face, body, stubby, and spiky networks in the VTC-SOM. First, based on the single-unit data, we calculated the average responses of neurons within the face, body, stubby, and spiky patches of the monkey ITC and identified ten most-preferred and ten non-preferred stimuli for each of the four categories. The selected stimuli were subsequently used to localize the face, body, spiky, and stubby network in the VTC-SOM. Specifically, for each category, we obtained the two average activation maps of the VTC-SOM for the most-preferred and non-preferred stimuli, and calculated a difference map between them. To match the number of functional regions revealed in the monkey ITC, we used a stricter criterion than that in Experiment 1 to define the category-selective regions. Here, for each category, units that showed the biggest difference (here, top 10 percent of all units) were defined as the category-selective units. In this way, we obtained the face, body, stubby, and spiky networks in the VTC-SOM.

Then, we calculated an RSM for each of the face, body, stubby, and spiky networks in the VTC-SOM. For each network, elements in its model RSM represented the measure of 1 - d (normalized Euclidean distance between the activation patterns of units elicited by two stimuli). The neural RSMs for the face, body, stubby, and spiky patches in the monkey ITC were obtained in the same way.

Finally, for each network, we computed a Pearson correlation coefficient between the VTC-SOM RSM and neural RSM of the monkey ITC (Figure 5).

### Testing the hierarchy of increasingly view-invariant representations in the face network of the VTC-SOM

To explore whether the hierarchy in view invariance found in Bao et al. (2020) was also present in our VTC-SOM, we took two face stimuli and two non-face stimuli (airplane and vase) from the stimulus-set for single-unit recording, each of which contained 8 different views, as examples to explore the view-invariant representation in three face-selective regions of the face network in the VTC-SOM. First, for each stimulus view, we obtained the activation pattern of units in each region of the face network in the VTC-SOM. Then, we computed an RSM (8×8) for each region of the face network. Each element in the RSM represented the correlation coefficient between activation patterns elicited by two views of the same stimulus for each region (Figure 6).

Finally, for each stimulus, we conducted the Kolmogorov–Smirnov test to compare the RSMs of two regions to examine whether the face network of the VTC-SOM contained a hierarchy of increasingly view-invariant representations.

## References

Avants, B. B., Epstein, C. L., Grossman, M., & Gee, J. C. (2008). Symmetric diffeomorphic image registration with cross-correlation: evaluating automated labeling of elderly and neurodegenerative brain. Medical image analysis, 12(1), 26–41.

Bao, P., She, L., McGill, M., & Tsao, D. Y. (2020). A map of object space in primate inferotemporal cortex. Nature, 583(7814), 103–108. doi:10.1038/s41586-020-2350-5

Beauchamp, M. S., Lee, K. E., Haxby, J. V., & Martin, A. (2002). Parallel visual motion processing streams for manipulable objects and human movements. Neuron, 34(1), 149–159.

Bednar, J. A., & Miikkulainen, R. (2003). Self-organization of spatiotemporal receptive fields and laterally connected direction and orientation maps. Neurocomputing, 52, 473–480.

Benavides-Piccione, R., Ballesteros-Yáñez, I., DeFelipe, J., & Yuste, R. (2002). Cortical area and species differences in dendritic spine morphology. Journal of neurocytology, 31(3), 337–346.

Chklovskii, D. B., & Koulakov, A. A. (2004). Maps in the brain: what can we learn from them? Annu Rev Neurosci, 27, 369–392. doi:10.1146/annurev.neuro.27.070203.144226

Cichy, R. M., Khosla, A., Pantazis, D., Torralba, A., & Oliva, A. (2016). Comparison of deep neural networks to spatio-temporal cortical dynamics of human visual object recognition reveals hierarchical correspondence. Sci Rep, 6, 27755. doi:10.1038/srep27755

DiCarlo, J. J., & Cox, D. D. (2007). Untangling invariant object recognition. Trends Cogn Sci, 11(8), 333–341. doi:10.1016/j.tics.2007.06.010

DiCarlo, J. J., Zoccolan, D., & Rust, N. C. (2012). How does the brain solve visual object recognition? Neuron, 73(3), 415–434.

Downing, P. E., Jiang, Y., Shuman, M., & Kanwisher, N. (2001). A cortical area selective for visual processing of the human body. Science, 293(5539), 2470–2473.

Durbin, R., & Mitchison, G. (1990). A dimension reduction framework for understanding cortical maps. Nature, 343(6259), 644–647.

Durbin, R., & Willshaw, D. (1987). An analogue approach to the travelling salesman problem using an elastic net method. Nature, 326(6114), 689–691.

Epstein, R., & Kanwisher, N. (1998). A cortical representation of the local visual environment. Nature, 392(6676), 598–601.

Erickson, C. A., & Desimone, R. (1999). Responses of macaque perirhinal neurons during and after visual stimulus association learning. Journal of Neuroscience, 19(23), 10404–10416.

Felleman, D. J., & Van Essen, D. C. (1991). Distributed hierarchical processing in the primate cerebral cortex. Cerebral cortex (New York, NY: 1991), 1(1), 1–47.

Gilbert, C. D., & Wiesel, T. N. (1983). Clustered intrinsic connections in cat visual cortex. Journal of Neuroscience, 3(5), 1116–1133.

Glasser, M. F., Coalson, T. S., Robinson, E. C., Hacker, C. D., Harwell, J., Yacoub, E., … Jenkinson, M. (2016). A multi-modal parcellation of human cerebral cortex. Nature, 536(7615), 171–178.

Grill-Spector, K., & Malach, R. (2004). The human visual cortex. Annu. Rev. Neurosci., 27, 649–677.

Grill-Spector, K., & Weiner, K. S. (2014). The functional architecture of the ventral temporal cortex and its role in categorization. Nature Reviews Neuroscience, 15(8), 536–548.

Grinvald, A., Lieke, E. E., Frostig, R. D., & Hildesheim, R. (1994). Cortical point-spread function and long-range lateral interactions revealed by real-time optical imaging of macaque monkey primary visual cortex. Journal of Neuroscience, 14(5), 2545–2568.

Guclu, U., & van Gerven, M. A. (2015). Deep Neural Networks Reveal a Gradient in the Complexity of Neural Representations across the Ventral Stream. J Neurosci, 35(27), 10005–10014. doi:10.1523/JNEUROSCI.5023-14.2015

Hasson, U., Avidan, G., Deouell, L. Y., Bentin, S., & Malach, R. (2003). Face-selective activation in a congenital prosopagnosic subject. Journal of cognitive neuroscience, 15(3), 419–431.

Hasson, U., Levy, I., Behrmann, M., Hendler, T., & Malach, R. (2002). Eccentricity bias as an organizing principle for human high-order object areas. Neuron, 34(3), 479–490.

Haxby, J. V., Gobbini, M. I., Furey, M. L., Ishai, A., Schouten, J. L., & Pietrini, P. (2001). Distributed and overlapping representations of faces and objects in ventral temporal cortex. Science, 293(5539), 2425–2430.

Haxby, J. V., Guntupalli, J. S., Connolly, A. C., Halchenko, Y. O., Conroy, B. R., Gobbini, M. I., … Ramadge, P. J. (2011). A common, high-dimensional model of the representational space in human ventral temporal cortex. Neuron, 72(2), 404–416.

Hebart, M. N., Zheng, C. Y., Pereira, F., & Baker, C. I. (2020). Revealing the multidimensional mental representations of natural objects underlying human similarity judgements. Nat Hum Behav, 4(11), 1173–1185. doi:10.1038/s41562-020-00951-3

Hegdé, J., & Van Essen, D. C. (2007). A comparative study of shape representation in macaque visual areas V2 and V4. Cerebral cortex, 17(5), 1100–1116.

Kanwisher, N., McDermott, J., & Chun, M. M. (1997). The fusiform face area: a module in human extrastriate cortex specialized for face perception. Journal of Neuroscience, 17(11), 4302–4311.

Khaligh-Razavi, S. M., & Kriegeskorte, N. (2014). Deep supervised, but not unsupervised, models may explain IT cortical representation. PLoS Comput Biol, 10(11), e1003915. doi:10.1371/journal.pcbi.1003915

Kobatake, E., & Tanaka, K. (1994). Neuronal selectivities to complex object features in the ventral visual pathway of the macaque cerebral cortex. Journal of neurophysiology, 71(3), 856–867.

Kohonen, T. (1989). Self-organizing feature maps. In Self-organization and associative memory (pp. 119–157): Springer.

Konkle, T. (2021). Emergent organization of multiple visuotopic maps without a feature hierarchy. bioRxiv.

Konkle, T., & Caramazza, A. (2013). Tripartite organization of the ventral stream by animacy and object size. J Neurosci, 33(25), 10235–10242. doi:10.1523/JNEUROSCI.0983-13.2013

Konkle, T., & Oliva, A. (2012). A real-world size organization of object responses in occipitotemporal cortex. Neuron, 74(6), 1114–1124.

Koulakov, A. A., & Chklovskii, D. B. (2001). Orientation preference patterns in mammalian visual cortex: a wire length minimization approach. Neuron, 29(2), 519–527.

Kriegeskorte, N., Mur, M., & Bandettini, P. A. (2008). Representational similarity analysis-connecting the branches of systems neuroscience. Frontiers in systems neuroscience, 2, 4.

Kriegeskorte, N., Mur, M., Ruff, D. A., Kiani, R., Bodurka, J., Esteky, H., … Bandettini, P. A. (2008). Matching categorical object representations in inferior temporal cortex of man and monkey. Neuron, 60(6), 1126–1141.

Krizhevsky, A. (2014). One weird trick for parallelizing convolutional neural networks. arXiv preprint 1404.5997.

Lee, H., Margalit, E., Jozwik, K. M., Cohen, M. A., Kanwisher, N., Yamins, D. L., & DiCarlo, J. J. (2020). Topographic deep artificial neural networks reproduce the hallmarks of the primate inferior temporal cortex face processing network. bioRxiv.

Levy, I., Hasson, U., Avidan, G., Hendler, T., & Malach, R. (2001). Center–periphery organization of human object areas. Nature neuroscience, 4(5), 533–539.

Linsker, R. (1988). Self-organization in a perceptual network. Computer, 21(3), 105–117.

Liu, X., Zhen, Z., & Liu, J. (2020). Hierarchical Sparse Coding of Objects in Deep Convolutional Neural Networks. Front Comput Neurosci, 14, 578158. doi:10.3389/fncom.2020.578158

Livingstone, M., & Hubel, D. (1988). Segregation of form, color, movement, and depth: anatomy, physiology, and perception. Science, 240(4853), 740–749.

Logothetis, N. K., & Sheinberg, D. L. (1996). Visual object recognition. Annual review of neuroscience, 19(1), 577–621.

Love, B. C., & Roads, B. D. (2021). Similarity as a Window on the Dimensions of Object Representation. Trends Cogn Sci, 25(2), 94–96. doi:10.1016/j.tics.2020.12.003

Malach, R., Levy, I., & Hasson, U. (2002). The topography of high-order human object areas. Trends in cognitive sciences, 6(4), 176–184.

Marr, D. (1982). Vision: A computational investigation into the human representation and processing of visual information, henry holt and co. Inc., New York, NY, 2(4.2).

McILWAIN, J. T. (1975). Visual receptive fields and their images in superior colliculus of the cat. Journal of neurophysiology, 38(2), 219–230.

Messinger, A., Squire, L. R., Zola, S. M., & Albright, T. D. (2001). Neuronal representations of stimulus associations develop in the temporal lobe during learning. Proc Natl Acad Sci U S A, 98(21), 12239–12244. doi:10.1073/pnas.211431098

Mishkin, M., Ungerleider, L. G., & Macko, K. A. (1983). Object vision and spatial vision: two cortical pathways. Trends in neurosciences, 6, 414–417.

Mohan, H., Verhoog, M. B., Doreswamy, K. K., Eyal, G., Aardse, R., Lodder, B. N., … Groot, C. (2015). Dendritic and axonal architecture of individual pyramidal neurons across layers of adult human neocortex. Cerebral cortex, 25(12), 4839–4853.

Nasr, S., Liu, N., Devaney, K. J., Yue, X., Rajimehr, R., Ungerleider, L. G., & Tootell, R. B. (2011). Scene-selective cortical regions in human and nonhuman primates. Journal of Neuroscience, 31(39), 13771–13785.

Nosofsky, R. M., Sanders, C. A., Meagher, B. J., & Douglas, B. J. (2018). Toward the development of a feature-space representation for a complex natural category domain. Behav Res Methods, 50(2), 530–556. doi:10.3758/s13428-017-0884-8

Obermayer, K., Blasdel, G. G., & Schulten, K. (1992). Statistical-mechanical analysis of self-organization and pattern formation during the development of visual maps. Physical Review A, 45(10), 7568.

Op de Beeck, H. P., Haushofer, J., & Kanwisher, N. G. (2008). Interpreting fMRI data: maps, modules and dimensions. Nat Rev Neurosci, 9(2), 123–135. doi:10.1038/nrn2314

Park, S. H., Cha, K., & Lee, S.-H. (2013). Coaxial anisotropy of cortical point spread in human visual areas. Journal of Neuroscience, 33(3), 1143–1156.

Riesenhuber, M. (2020). How the mind sees the world. Nat Hum Behav, 4(11), 1100–1101. doi:10.1038/s41562-020-00973-x

Sakai, K., & Miyashita, Y. (1991). Neural organization for the long-term memory of paired associates. Nature, 354(6349), 152–155.

Shiffrin, R. M., Bassett, D. S., Kriegeskorte, N., & Tenenbaum, J. B. (2020). The brain produces mind by modeling. Proc Natl Acad Sci U S A, 117(47), 29299–29301. doi:10.1073/pnas.1912340117

Van Essen, D. C., Smith, S. M., Barch, D. M., Behrens, T. E., Yacoub, E., Ugurbil, K., & Consortium, W.-M. H. (2013). The WU-Minn human connectome project: an overview. Neuroimage, 80, 62–79.

Warrington, E. K., & McCarthy, R. (1983). Category specific access dysphasia. Brain, 106(4), 859–878.

Weigand, M., Sartori, F., & Cuntz, H. (2017). Universal transition from unstructured to structured neural maps. Proc Natl Acad Sci U S A, 114(20), E4057–E4064. doi:10.1073/pnas.1616163114

Weiner, K. S., & Grill-Spector, K. (2013). Neural representations of faces and limbs neighbor in human high-level visual cortex: evidence for a new organization principle. Psychological research, 77(1), 74–97.

Wen, H., Shi, J., Zhang, Y., Lu, K. H., Cao, J., & Liu, Z. (2018). Neural Encoding and Decoding with Deep Learning for Dynamic Natural Vision. Cereb Cortex, 28(12), 4136–4160. doi:10.1093/cercor/bhx268

Yamins, D. L., Hong, H., Cadieu, C. F., Solomon, E. A., Seibert, D., & DiCarlo, J. J. (2014). Performance-optimized hierarchical models predict neural responses in higher visual cortex. Proceedings of the National Academy of Sciences, 111(23), 8619–8624.

